## Supplementary Information for "Wildflower phenological escape differs by continent and spring temperature"

### Supplementary Methods

### Model design and spatial autocorrelation

*Summary of INLA spatial autocorrelation analysis*

There were no significant differences (and high similarity in mean values) among the estimated intercepts or phenological sensitivities to spring temperature and elevation when comparing the posterior estimates from the analogous JAGS and INLA-SPDE models (Table S4). This suggests that the high fidelity between the two approaches found by other studies apply to our phenological models as well.

In contrast, incorporating spatial autocorrelation into the model structure changed model fit and performance as well as posterior estimates of parameter values, sometimes in statistically significant ways (Tables S3-S4). The model versions that included spatial autocorrelation had elevated predicted vs. observed $r^2$ values and significantly lower DIC values compared to those that did not for all combinations of plant group and continent (Table S3). Specifically, models that included spring temperature and elevation as fixed effects, species as a random effect, and the spatial autocorrelation term were the best-performing or among the best-performing model structures for five of the six plant group-continent combinations (the only exception being tree LOD in Asia, which was better predicted by either including winter temperature or annual precipitation instead of elevation).

Including spatial autocorrelation had a particularly strong effect on model fit and performance in Asia and Europe (Table S3), where predicted vs. observed $r^2$ values increased by 69-86% and 50-65%, respectively. North American models, which had the best-performing models prior to the addition of spatial autocorrelation terms, only experienced increases in predicted vs. observed $r^2$ values of 3-6%. In contrast to these large differences in how model performance was affected, the only parameter values that significantly changed with the addition of spatial autocorrelation terms were in the North American models (Table S4). North American phenological sensitivity to spring temperature was significantly less negative with the addition of spatial autocorrelation terms for both wildflower FFD and canopy tree LOD models. Importantly, the difference in phenological sensitivity between trees and wildflowers that was statistically significant in the original model structure disappeared – there was no significant difference in sensitivity to spring temperature when spatial autocorrelation was included in the model. This was the only continent where the relationship in spring temperature sensitivity between wildflowers and canopy trees changed in response to adding spatial autocorrelation into the model. Model intercepts and sensitivities to elevation did not significantly change for any plant group-continent combination when spatial autocorrelation was added to the model structure (Table S4).

*Justification for exclusion of spatial autocorrelation*

The addition of the spatial autocorrelation term had qualitatively different effects depending on the continent. This is most noticeable for the Asian models, where the posterior-predicted mean effects of the spatial autocorrelation term ranged from -37.2 to 50.1 days for trees and -38.8 to 36.0 days for wildflowers (Fig. S2), meaning that the spatial relationship between data points altered predicted phenology by more than a month in either direction even after spring temperature and elevation had been accounted for in the model structure. This range was narrower in both Europe (Fig. S3; -10.8 to 6.0 days for trees and -24.6 to 33.1 days for wildflowers) and eastern North America (Fig. S4, -18.8 to 19.3 days for trees and -10.3 to 9.5 days for wildflowers), suggesting that spatial relationships (which could include relationships with climate or geographic drivers that covary with the spatial relationships) are more influential on the predictive accuracy of spring phenology models in Asia compared to on the other two continents.

Potentially even more apparent when comparing Figures S2-S4 is the difference in the smooth clines in North America and the patchy, hot-spot patterns in Europe and Asia. The patterns in North America appear to follow latitudinal trends, potentially reflecting photoperiodic effects that have been previously found to strongly affect spring phenology of deciduous tree species[1]. Europe is somewhat like North America in that there are relatively smooth gradients that appear to be somewhat linked to geographic features (for example, strong spatial autocorrelative effects appear for wildflower species along major mountain ranges). However, different ranges have different effects on phenology with delayed flowering in the Pyrenees and earlier flowering in the Alps. On one hand, this makes some sense because our models already account for elevational effects, but on the other hand, it seems to contradict other potential drivers associated with mountains such as the timing of snowmelt. It is also interesting to note that our results qualitatively differ from a recent European study also using herbarium collections to model spring phenology shifts[2], which found early-shifts associated with both mountain ranges when elevation, year, spring and winter temperature, and spring precipitation had all been accounted for as phenological drivers.

Sharply contrasting the North American and European results, however, spatial autocorrelation in East Asia appears extremely heterogeneous, with regions of extremely delayed phenology immediately next to regions with strong early shifts (Fig. S2). We could think of no reasonable explanations to explain the observed patterns of spatial autocorrelative effects (for example, some adjacent areas differed by a total of about 2 months in predicted effects) and therefore posit that they are the result of nuances in where and how the herbarium specimens were collected. In support of this hypothesis, Asian spatial autocorrelation had overall much wider standard deviation (Fig. S2b, d) compared to both Europe and North America, suggesting a greater degree of uncertainty in the spatial relationships inherent to the collections. This appears to be, in part, due to the concentration of collections in certain areas (i.e., the areas with much lower standard deviations). That is, areas with strong spatial autocorrelational mean effects are typically also the areas with the lowest standard deviation, suggesting that collections are concentrated in these areas. This could therefore also suggest that some regions were simply collected earlier on average than others as a function of logistical constraints (e.g., it is easier to collect specimens closer to urban areas as opposed to rural areas) rather than through accurate reflection of ecological processes.

Due to the inconsistencies in how spatial autocorrelation affected predicted phenology across the three continents, we decided not to include this term in the models presented in the main body of the manuscript. Still, we want to highlight the importance in how these effects can influence model performance both in terms of predicted effects (e.g., spring temperature sensitivity changed for North American plant species depending on if spatial autocorrelation was included in the model or not) and in terms of predictive power (European and Asian models experienced substantial increases in predicted vs. observed $r^2$ values when spatial autocorrelation was added to the models). We would also like to note that because the nuances and discrepancies that arose with the Asian phenology models are likely related to concentrated observations and uneven sampling distributions, further collections and digitization efforts should resolve these biases in the near future.

### Supplementary Tables

**Table S1:** Number of individual specimens of each species used in this study. Total number of specimens across all continents N = 5,522.

| Region | Lifeform | Species | #Specimens |
|---|---|---|---|
| Eastern Asia | Trees n = 899 | *Acer pictum subsp. Mono* (Maxim.) | 37 |
| | | *Acer truncatum* (Bunge) | 94 |
| | | *Betula platyphylla* (Sukaczev) | 113 |
| | | *Fraxinus chinensis* (Roxb.) | 99 |
| | | *Juglans mandshurica* (Maxim.) | 53 |
| | | *Populus davidiana* (Dode) | 125 |
| | | *Quercus aliena* (Blume) | 44 |
| | | *Quercus mongolica* (Fisch.) | 66 |
| | | *Quercus variabilis* (Blume) | 186 |
| | | *Ulmus pumila* (L.) | 82 |
| | Wildflowers n = 1,418 | *Androsace umbellata* (Lour.) | 486 |
| | | *Astragalus scaberrimus* (Bunge) | 203 |
| | | *Gueldenstaedtia verna* (Georgi) | 185 |
| | | *Leibnitzia anandria* (L.) | 206 |
| | | *Pulsatilla chinensis* (Bunge) | 177 |
| | | *Viola prionantha* (Bunge) | 161 |
| Europe | Trees n = 532 | *Acer platanoides* (L.) | 96 |
| | | *Acer pseudoplatanus* (L.) | 69 |
| | | *Carpinus betulus* (L.) | 62 |
| | | *Fagus sylvatica* (L.) | 184 |
| | | *Quercus petraea* (Matt.) | 54 |
| | | *Quercus robur* (L.) | 67 |
| | Wildflowers n = 618 | *Allium ursinum* (L.) | 79 |
| | | *Anemone nemorosa* (L.) | 130 |
| | | *Anemone ranunculoides* (L.) | 52 |
| | | *Corydalis cava* (L.) | 124 |
| | | *Corydalis solida* (L.) | 50 |
| | | *Hepatica nobilis* (L.) | 183 |
| Eastern North America | Trees n = 995 | *Acer rubrum* (L.) | 308 |
| | | *Acer saccharum* (Marshall) | 138 |
| | | *Carya glabra* (Miller) | 111 |
| | | *Fagus grandifolia* (Ehrh.) | 161 |
| | | *Quercus alba* (L.) | 158 |
| | | *Quercus rubra* (L.) | 119 |
| | Wildflowers n = 1,060 | *Anemone quinquefolia* (L.) | 188 |
| | | *Dicentra canadensis* (L.) | 143 |
| | | *Dicentra cucullaria* (L.) | 169 |
| | | *Erythronium americanum* (Ker-Gawl.) | 191 |
| | | *Hepatica americana* (DC.) | 148 |
| | | *Sanguinaria canadensis* (L.) | 221 |

**<u>Table S2:</u>** Means and standard deviations of day of year of observed phenology and estimated spring temperature of each observation, grouped by continent and lifeform (trees or wildflowers).

| Continent | Group | Observed Day of Year | | March-April Temperature (°C) | |
|---|---|---|---|---|---|
| | | Mean | S.d. | Mean | S.d. |
| Asia | Trees | 115.72 | 24.83 | 9.12 | 4.57 |
| | Wildflowers | 116.77 | 24.44 | 8.48 | 4.32 |
| Europe | Trees | 132.24 | 15.12 | 5.54 | 2.44 |
| | Wildflowers | 115.92 | 20.75 | 5.01 | 2.80 |
| N. America | Trees | 116.87 | 22.57 | 8.84 | 5.42 |
| | Widlflowers | 110.55 | 19.00 | 7.19 | 4.52 |

**<u>Table S3:</u>** Model comparison statistics for the different model versions evaluated as part of this study. Values in bold italics indicate the model(s) with the best deviance information criterion (DIC) value, assessed as being at least two less than all other models for the combination of continent and plant lifeform (wildflower or tree).

| Region | Model | Wildflower FFD | | Canopy Tree LOD | |
|---|---|---|---|---|---|
| | | $r^2$ | DIC | $r^2$ | DIC |
| Asia | Spring T | 0.388 | 12407.66 | 0.437 | 7832.79 |
| | SpringT + S.A. | ***0.740*** | ***11881.22*** | 0.738 | 7556.18 |
| | Spring T + Elevation | 0.398 | 12385.98 | 0.437 | 7833.75 |
| | Spring T + S.A. + Elevation | ***0.739*** | ***11881.01*** | 0.738 | 7557.06 |
| | Spring T + S.A. + Winter T | 0.739 | 11885.25 | ***0.740*** | ***7551.11*** |
| | Spring T + S.A. + Annual Prec. | 0.739 | 11886.79 | ***0.739*** | ***7551.04*** |
| | Spring T + S.A. + Spring Prec. | ***0.741*** | ***11882.06*** | 0.737 | 7557.83 |
| Europe | Spring T | 0.356 | 5246.99 | 0.236 | 4270.80 |
| | SpringT + S.A. | 0.678 | 5012.63 | 0.387 | 4248.59 |
| | Spring T + Elevation | 0.407 | 5198.44 | 0.252 | 4261.37 |
| | Spring T + S.A. + Elevation | ***0.672*** | ***5005.54*** | ***0.380*** | ***4237.75*** |
| | Spring T + S.A. + Winter T | 0.678 | 5013.31 | 0.370 | 4248.04 |
| | Spring T + S.A. + Annual Prec. | 0.679 | 5013.94 | 0.388 | 4248.09 |
| | Spring T + S.A. + Spring Prec. | 0.678 | 5014.48 | 0.382 | 4249.59 |
| North America | Spring T | 0.705 | 7968.63 | 0.754 | 7646.52 |
| | SpringT + S.A. | 0.725 | 7939.67 | 0.814 | 7498.66 |
| | Spring T + Elevation | 0.705 | 7969.54 | 0.761 | 7621.36 |
| | Spring T + S.A. + Elevation | ***0.726*** | ***7927.93*** | ***0.809*** | ***7483.74*** |
| | Spring T + S.A. + Winter T | 0.733 | 7943.83 | 0.814 | 7500.58 |
| | Spring T + S.A. + Annual Prec. | 0.731 | 7941.40 | 0.814 | 7500.18 |
| | Spring T + S.A. + Spring Prec. | 0.725 | 7939.06 | 0.814 | 7499.84 |

**Table S4:** Posterior estimated mean and 95% Bayesian credible intervals of model slopes and intercepts for models that include average spring temperature and elevation as fixed effects. Posterior estimates are presented for models both with and without spatial autocorrelation terms.

| | | | **Sp. Temp. + Elev. (JAGS)** | | | **Sp. Temp. + Elev. (INLA)** | | | **Sp. Temp., Elev., + S.A.** | | |
|---|---|---|---|---|---|---|---|---|---|---|---|
| | | | **2.5% CI** | **Mean** | **97.5% CI** | **2.5% CI** | **Mean** | **97.5% CI** | **2.5% CI** | **Mean** | **97.5% CI** |
| Asia | FFD | $\beta_1$ (SpT) | -3.364 | -3.118 | -2.861 | -3.382 | -3.131 | -2.879 | -3.417 | -3.053 | -2.686 |
| | | Intercept | 96.78 | 120.46 | 144.11 | 111.33 | 134.90 | 158.46 | 112.31 | 136.09 | 159.85 |
| | | $\beta_2$ (Elev) | 2.335 | 3.794 | 5.242 | 2.239 | 3.733 | 5.227 | 0.406 | 2.940 | 5.482 |
| | LOD | $\beta_1$ (SpT) | -2.971 | -2.621 | -2.261 | -3.000 | -2.647 | -2.293 | -2.834 | -2.354 | -1.865 |
| | | Intercept | 110.57 | 129.37 | 148.13 | 119.28 | 138.30 | 157.31 | 116.45 | 135.98 | 155.49 |
| | | $\beta_2$ (Elev) | -2.544 | -0.840 | 0.882 | -2.668 | -0.894 | 0.878 | -2.530 | 0.335 | 3.263 |
| Europe | FFD | $\beta_1$ (SpT) | -3.476 | -3.021 | -2.565 | -3.510 | -3.044 | -2.578 | -3.833 | -3.245 | -2.656 |
| | | Intercept | 85.64 | 108.91 | 132.54 | 99.49 | 123.11 | 146.70 | 103.35 | 127.29 | 151.21 |
| | | $\beta_2$ (Elev) | 10.722 | 14.768 | 18.678 | 10.694 | 14.682 | 18.666 | 6.940 | 12.538 | 18.103 |
| | LOD | $\beta_1$ (SpT) | -3.268 | -2.792 | -2.298 | -3.310 | -2.815 | -2.321 | -3.440 | -2.853 | -2.259 |
| | | Intercept | 101.39 | 125.22 | 149.10 | 116.32 | 139.94 | 163.55 | 115.51 | 139.31 | 163.08 |
| | | $\beta_2$ (Elev) | 3.094 | 7.098 | 11.162 | 2.997 | 7.093 | 11.187 | 4.293 | 9.649 | 15.297 |
| North America | FFD | $\beta_1$ (SpT) | -3.276 | -3.141 | -3.004 | -3.279 | -3.141 | -3.002 | -2.793 | -2.403 | -2.009 |
| | | Intercept | 91.11 | 113.95 | 136.98 | 104.66 | 128.16 | 151.63 | 91.06 | 119.70 | 147.39 |
| | | $\beta_2$ (Elev) | -1.394 | 1.779 | 4.951 | -1.467 | 1.715 | 4.894 | 2.854 | 7.189 | 11.565 |
| | LOD | $\beta_1$ (SpT) | -3.756 | -3.617 | -3.485 | -3.752 | -3.619 | -3.485 | -2.865 | -2.422 | -1.977 |
| | | Intercept | 104.03 | 127.07 | 149.88 | 117.90 | 141.40 | 164.89 | 105.11 | 131.68 | 157.58 |
| | | $\beta_2$ (Elev) | 6.863 | 10.835 | 15.037 | 6.733 | 10.765 | 14.793 | 8.502 | 14.257 | 20.009 |

**<u>Table S5:</u>** Posterior estimated species-level random effects (means and 95% Bayesian credible intervals) for East Asian tree and wildflower species. Estimates are provided for the parameters derived from the JAGS model described in the main methods section.

| | **Posterior estimates for JAGS model** | | |
|---|---|---|---|
| Tree Species | 2.5 BCI | Mean | 97.5 BCI |
| *Acer pictum* ssp. *mono* | -4.02 | 16.032 | 34.763 |
| *Acer truncatum* | -7.489 | 11.93 | 31.63 |
| *Betula platyphylla* | -3.439 | 16.033 | 34.84 |
| *Fraxinus chinensis* | -0.761 | 18.535 | 37.14 |
| *Juglans mandshurica* | 3.872 | 23.562 | 42.211 |
| *Populus davidiana* | -20.846 | -1.648 | 17.295 |
| *Quercus aliena* | -1.879 | 17.659 | 37.179 |
| *Quercus mongolica* | 4.582 | 24.411 | 43.4 |
| *Quercus variabilis* | -8.225 | 11.018 | 30.302 |
| *Ulmus pumila* | -33.017 | -12.825 | 6.226 |
| Wildflower Species | | | |
| *Androsace umbellata* | -2.874 | 20.763 | 44.233 |
| *Astragalus scaberrimus* | 1.773 | 25.941 | 49.428 |
| *Gueldenstaedtia verna* | -2.861 | 21.045 | 43.999 |
| *Leibnitzia anandria* | -0.886 | 22.688 | 46.489 |
| *Pulsatilla chinensis* | -6.085 | 17.687 | 41.292 |
| *Viola prionantha* | -11.351 | 12.456 | 35.209 |

**Table S6:** Posterior estimated species-level random effects (means and 95% Bayesian credible intervals) for European tree and wildflower species. Estimates are provided for the parameters derived from the JAGS model described in the main methods section.

| | **Posterior estimates for JAGS model** | | |
|---|---|---|---|
| Tree Species | 2.5 BCI | Mean | 97.5 BCI |
| *Acer platanoides* | -10.918 | 12.977 | 36.649 |
| *Acer pseudoplatanus* | 1.272 | 25.373 | 49.333 |
| *Carpinus betulus* | -9.569 | 14.034 | 38.254 |
| *Fagus sylvatica* | -2.973 | 20.898 | 44.675 |
| *Quercus petraea* | 4.917 | 28.551 | 52.648 |
| *Quercus robur* | 1.771 | 25.543 | 49.512 |
| Wildflower Species | | | |
| *Allium ursinum* | 16.907 | 40.499 | 64.238 |
| *Anemone nemorosa* | -8.603 | 15.288 | 38.582 |
| *Anemone ranunculoides* | -5.891 | 17.459 | 40.899 |
| *Corydalis cava* | -10.094 | 13.842 | 37.366 |
| *Corydalis solida* | -18.657 | 5.05 | 29.134 |
| *Hepatica nobilis* | -8.689 | 15.07 | 38.049 |

**<u>Table S7:</u>** Posterior estimated species-level random effects (means and 95% Bayesian credible intervals) for Eastern North American tree and wildflower species. Estimates are provided for the parameters derived from the JAGS model described in the main methods section.

| | **Posterior estimates for JAGS model** | | |
|---|---|---|---|
| Tree Species | 2.5 BCI | Mean | 97.5 BCI |
| *Acer rubrum* | -7.604 | 14.973 | 37.502 |
| *Acer saccharum* | -6.973 | 15.713 | 38.429 |
| *Carya glabra* | 7.179 | 30.13 | 52.733 |
| *Fagus grandifolia* | -4.443 | 18.569 | 41.082 |
| *Quercus alba* | 3.722 | 26.84 | 49.571 |
| *Quercus rubra* | -1.691 | 20.884 | 43.75 |
| Wildflower Species | | | |
| *Anemone quinquefolia* | 5.866 | 28.665 | 51.847 |
| *Dicentra canadensis* | 0.89 | 23.615 | 46.774 |
| *Dicentra cucullaria* | -3.67 | 19.123 | 42.092 |
| *Erythronium americanum* | -5.344 | 17.539 | 40.383 |
| *Hepatica americana* | -10.444 | 12.357 | 35.284 |
| *Sanguinaria canadensis* | -11.023 | 11.935 | 34.784 |

**<u>Supplementary Figures</u>**

**<u>Figure S1</u>**: Maps of the 5,522 herbarium specimens collected between 1901 and 2020 across a) eastern Asia, b) western Europe, and c) eastern North America. Points represent observations of wildflower FFD (circles) and overstory LOD (crosses). Background color indicates average March-April temperatures in the current climate simulation (averaged from 2009-2018 using CRU TS v4.03 climate data).

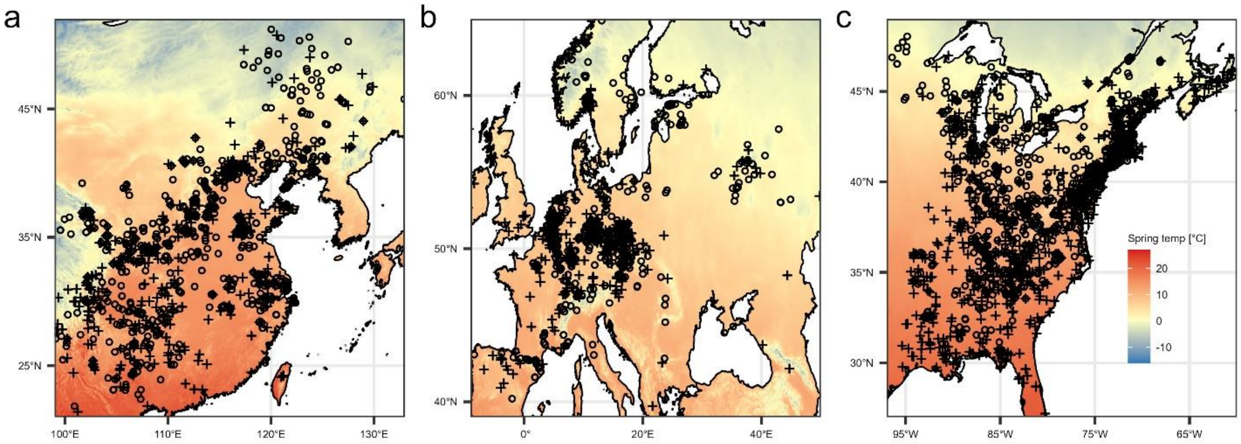

**Figure S2:** Posterior estimated means (left-hand panels) and standard deviations (right-hand panels) of spatial autocorrelation effects (in days) for (a-b) wildflower and (c-d) canopy tree phenology in East Asia. Positive and negative mean values indicate regions where phenology is expected to be delayed or earlier, respectively, compared to what is predicted from spring temperature and elevation alone. Maps are cropped to the extent of the Matérn meshes used to fit the models.

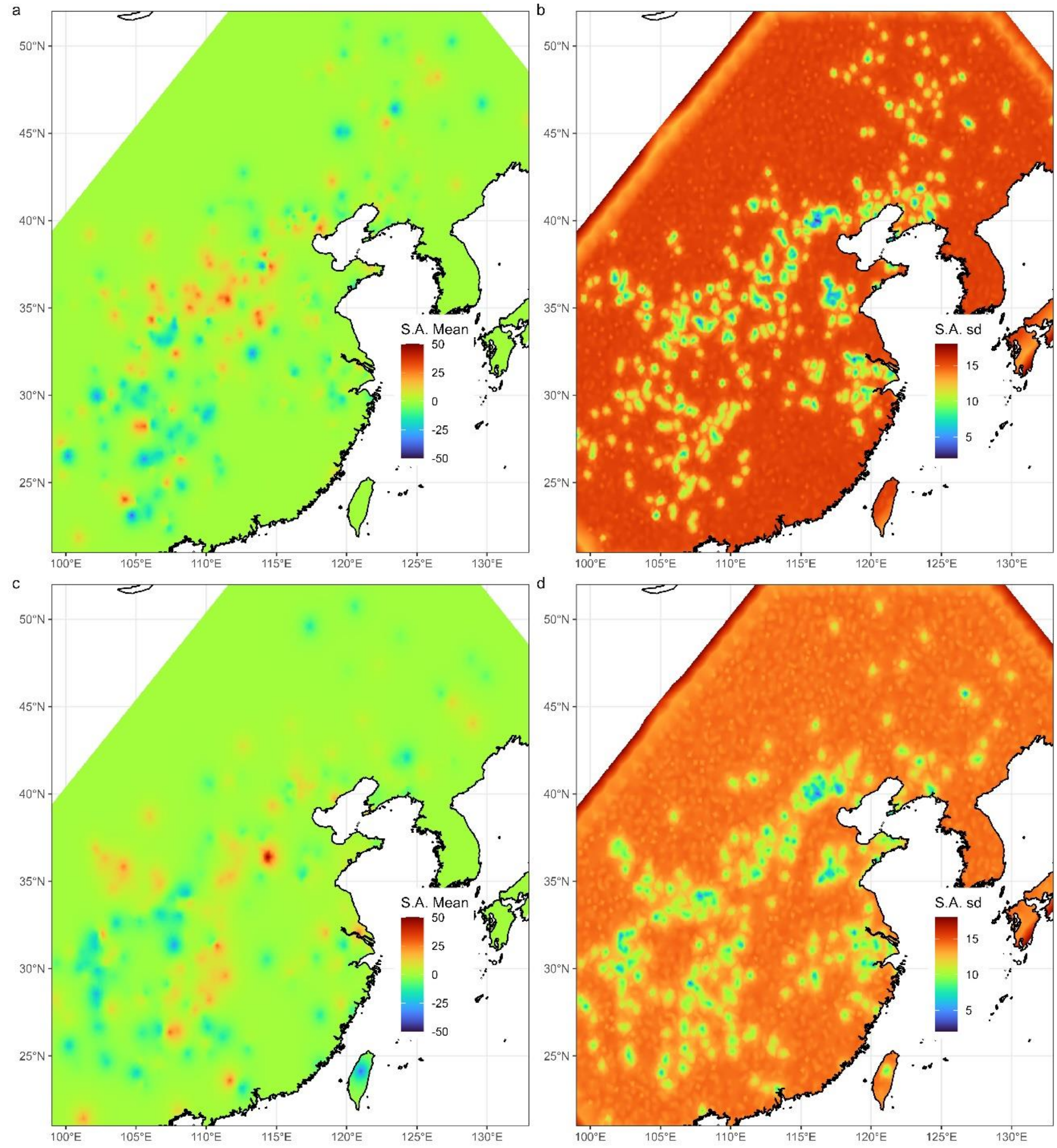

**Figure S3:** Posterior estimated means (left-hand panels) and standard deviations (right-hand panels) of spatial autocorrelation effects (in days) for (a-b) wildflower and (c-d) canopy tree phenology in Europe. Positive and negative mean values indicate regions where phenology is expected to be delayed or earlier, respectively, compared to what is predicted from spring temperature and elevation alone. Maps are cropped to the extent of the Matérn meshes used to fit the models.

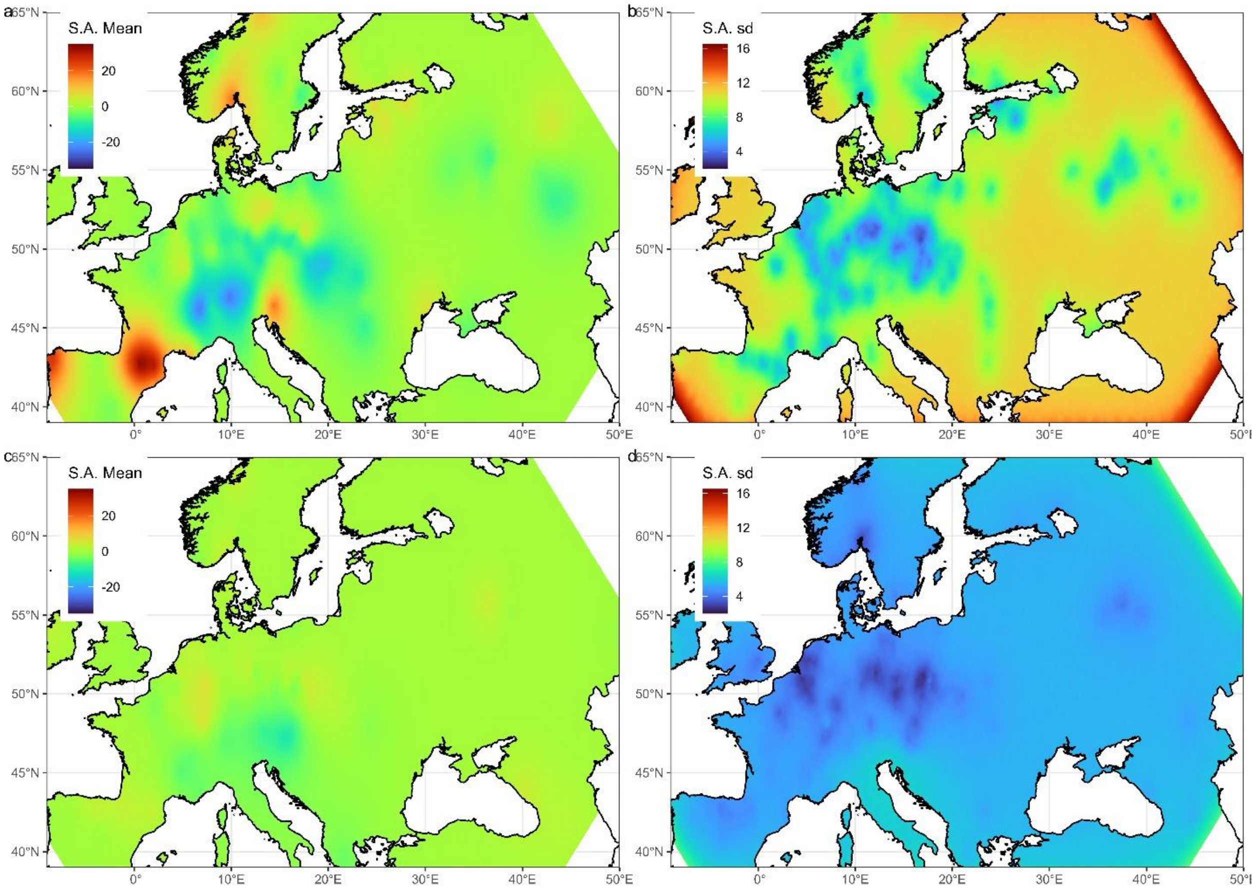

**Figure S4:** Posterior estimated means (left-hand panels) and standard deviations (right-hand panels) of spatial autocorrelation effects (in days) for (a-b) wildflower and (c-d) canopy tree phenology in eastern North America. Positive and negative mean values indicate regions where phenology is expected to be delayed or earlier, respectively, compared to what is predicted from spring temperature and elevation alone. Maps are cropped to the extent of the Matérn meshes used to fit the models.

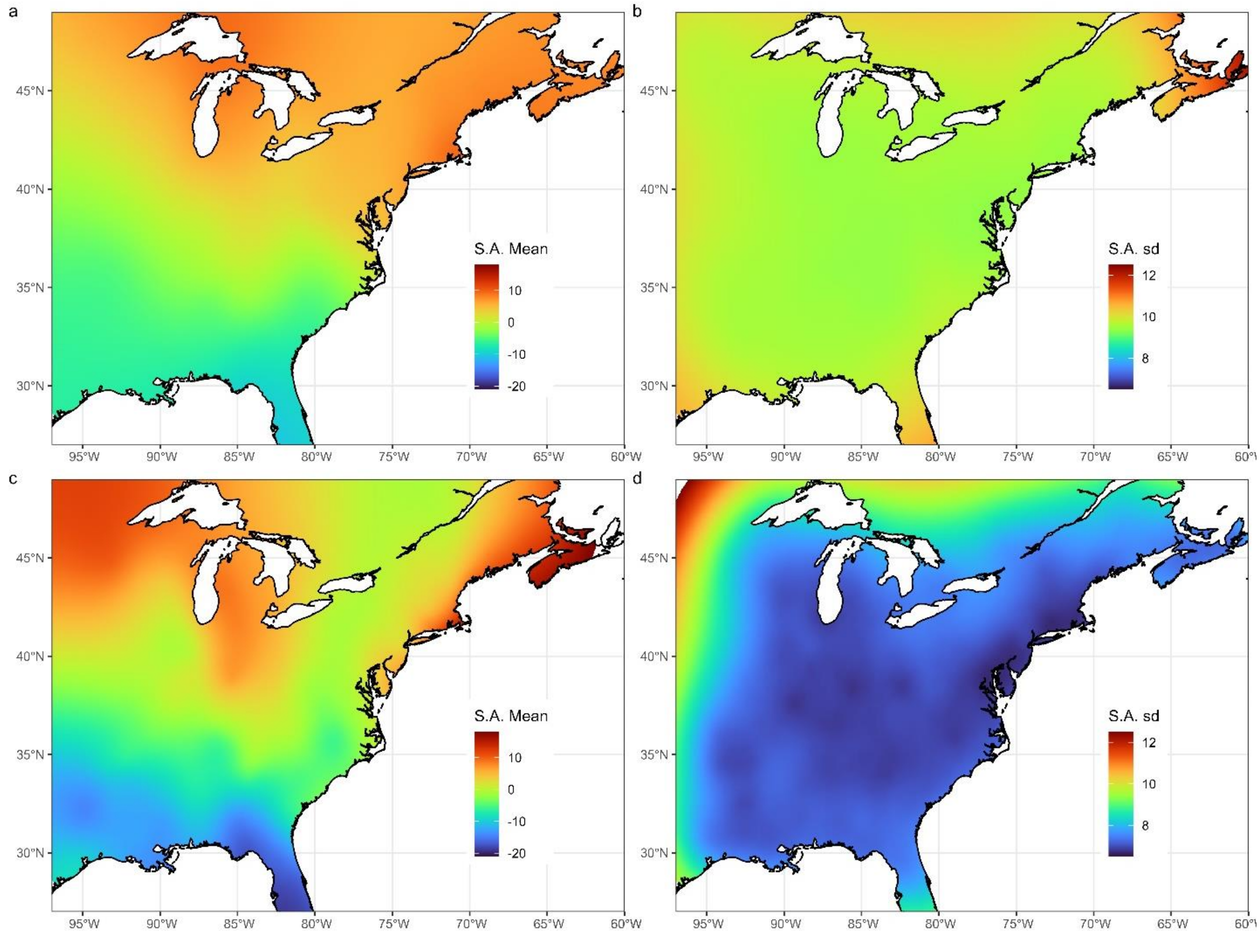

**Figure S5:** Estimated wildflower FFD (top row) and canopy tree LOD (bottom row) under current climate conditions (averaged from 2009-2018, see methods) in (a, d) Asia, (b, e) Europe, and c, f) North America.

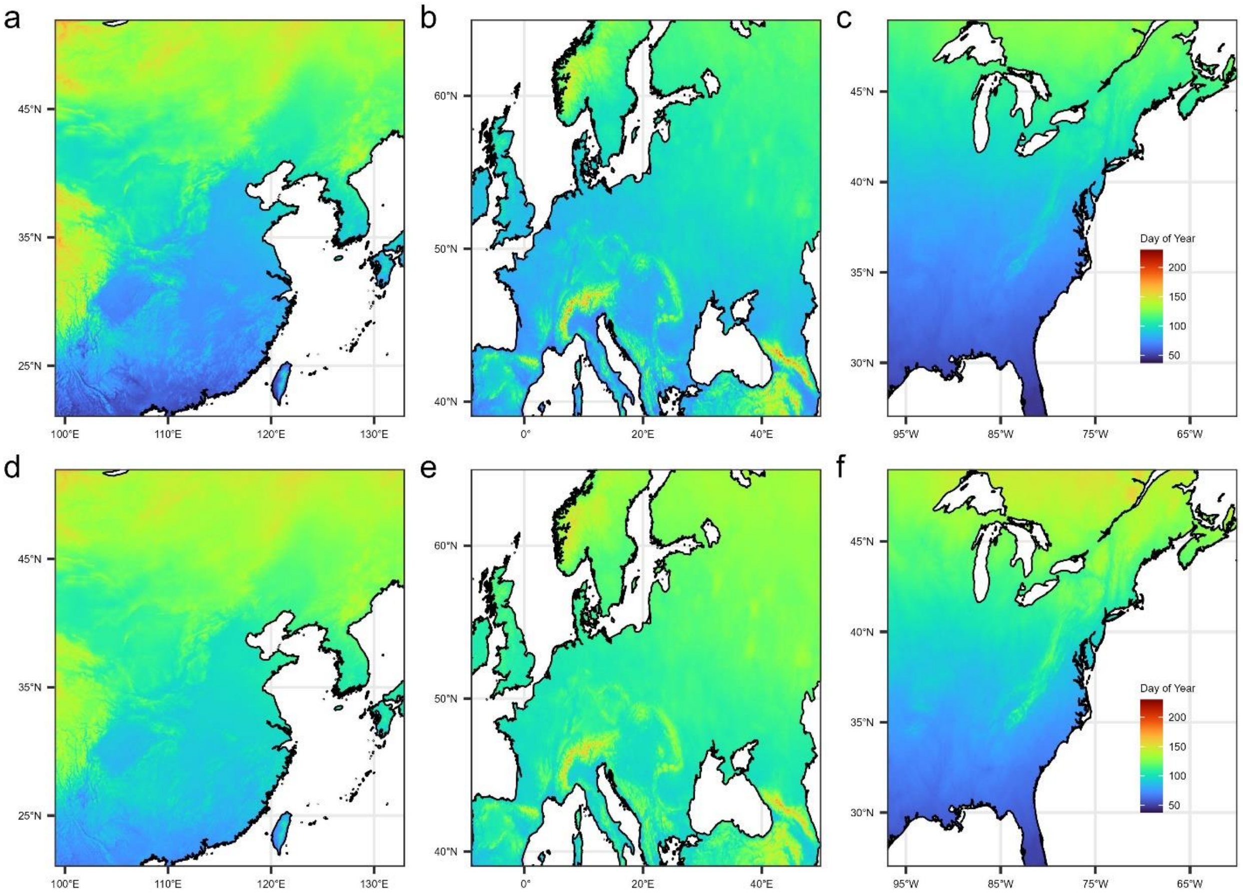

**Figure S6:** Projected wildflower FFD (top row) and canopy tree LOD (bottom row) under future climate conditions (estimated from the average projected climate conditions from 2081-2100, see methods) in (a, d) Asia, (b, e) Europe, and c, f) North America.

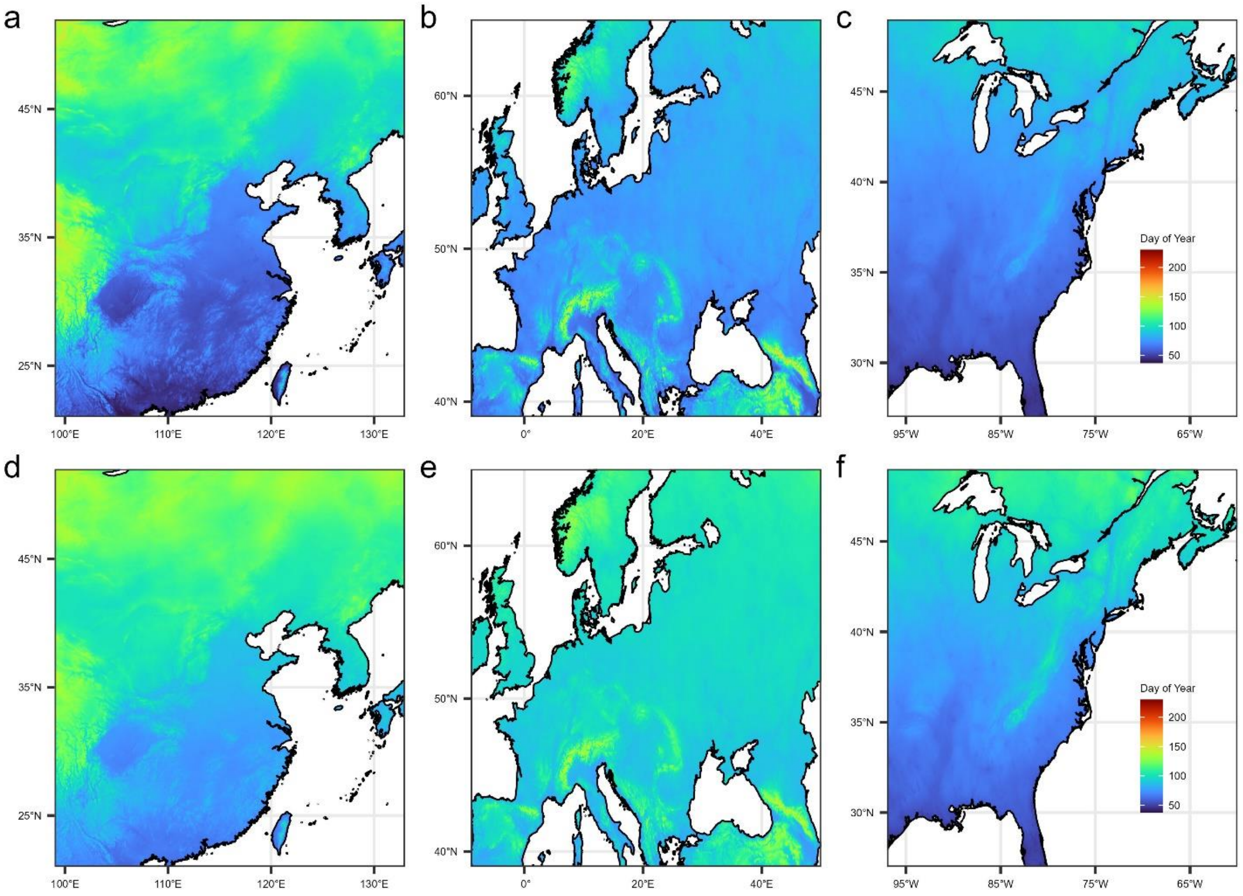

**Figure S7:** Projected mean difference between wildflower flowering (FFD) and canopy tree leaf out (LOD) (in days) under future climate conditions at the end of the century (see methods) in a) Asia, b) Europe, and c) North America. Negative values indicate LOD is estimated to occur before FFD.

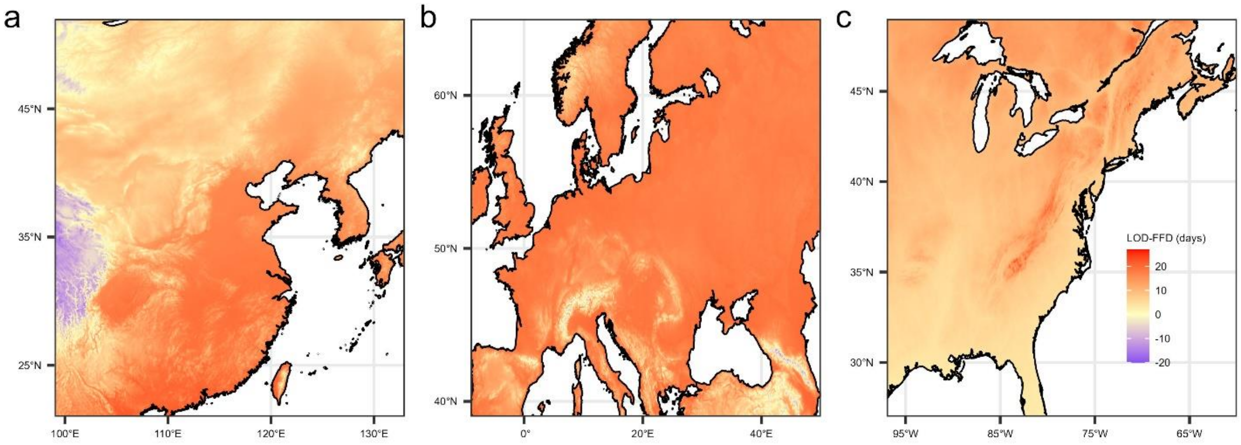

**Figure S8:** Difference in average spring temperature between current environmental conditions (averaged between 2009-2018) and future conditions (2080-2100) for a) Asia, b) Europe, and c) eastern North America. Dark gray regions indicate areas where the consensus land classification is < 1% deciduous or mixed deciduous forest cover.

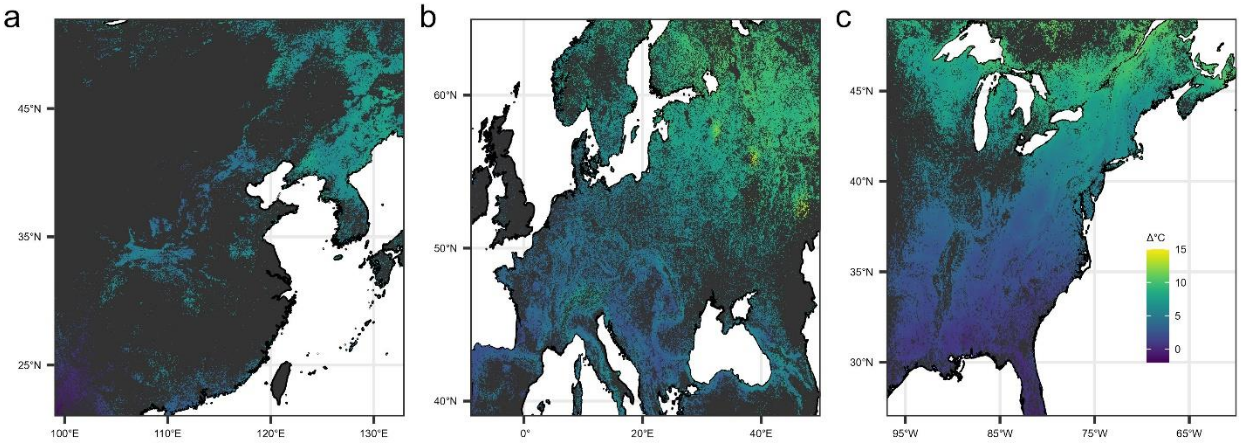

**Figure S9:** Correlation strengths (adjusted $r^2$ values) for linear regressions relating different one-, two-, and three-month aggregated spring temperature windows to observed wildflower First Flowering Date (FFD, blue) and tree Leaf Out Date (LOD, gold). Horizontal lines indicate the adjusted $r^2$ value for the relationship with average March-April temperature for ease of comparison with the other windows.

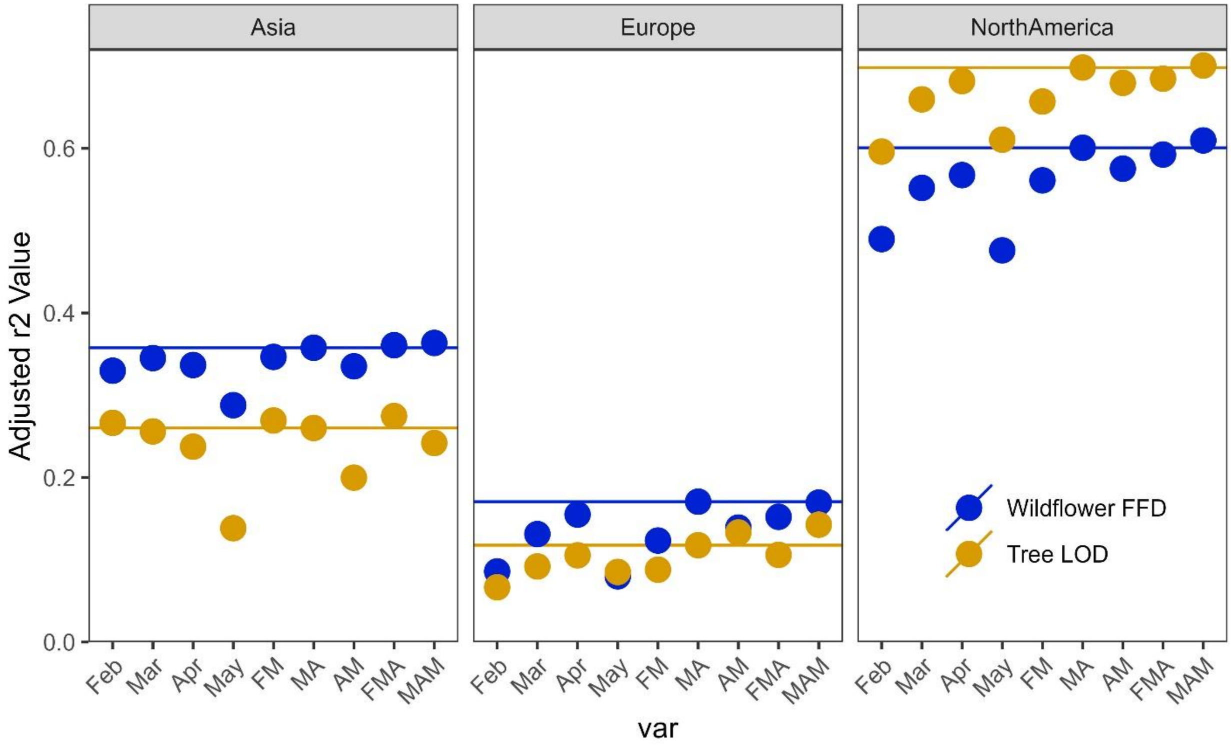

**Figure S10:** Matérn meshes used to fit INLA models that include spatial autocorrelation terms for a) East Asia, b) Europe, and c) Eastern North America. Red and blue points represent relative locations of observed wildflower and tree phenology, respectively. Coordinates are scaled from decimal degree coordinates so that all locations have positive latitude and longitude values.

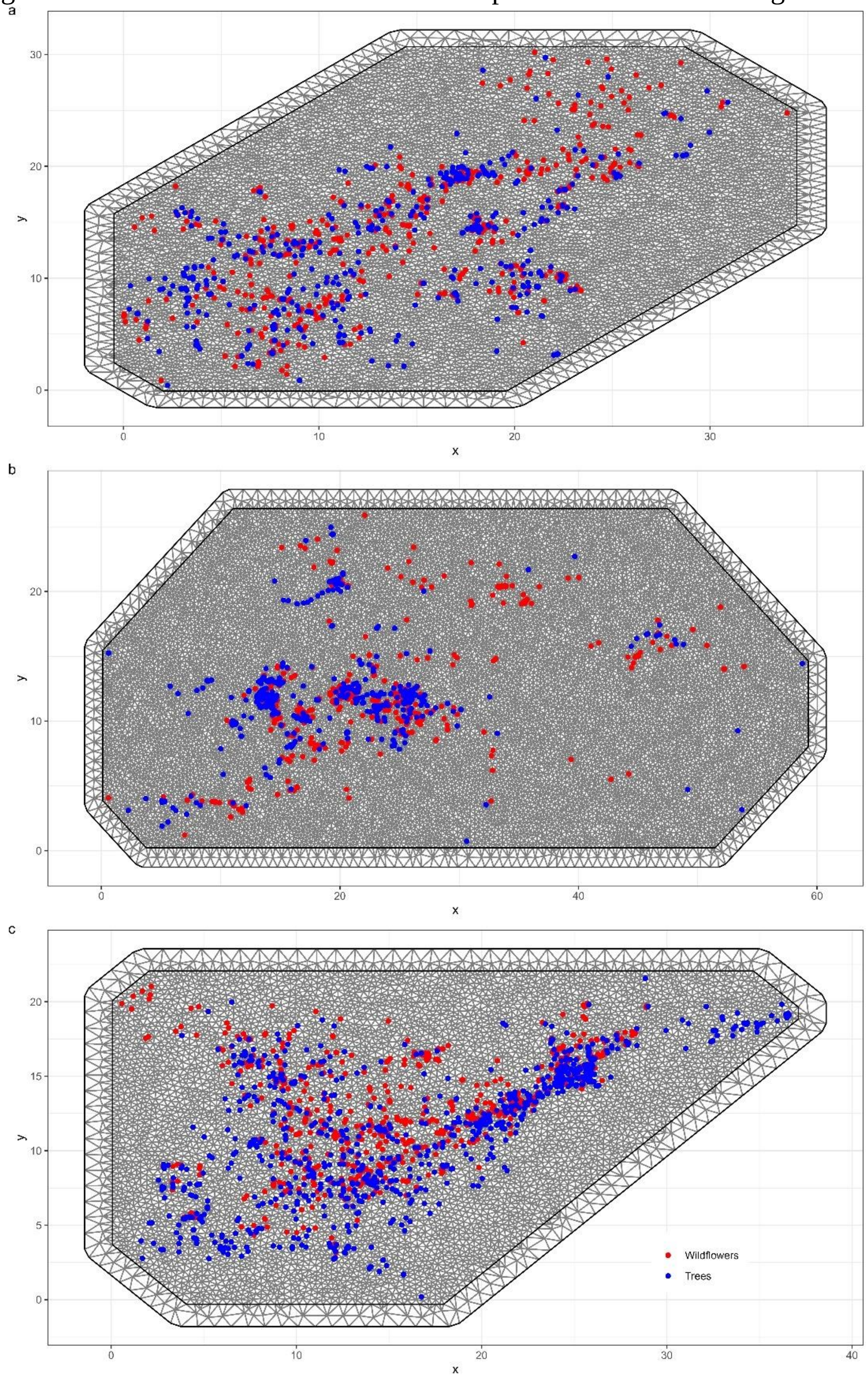

**Figure S11:** Full, uncropped version of Figure 3.

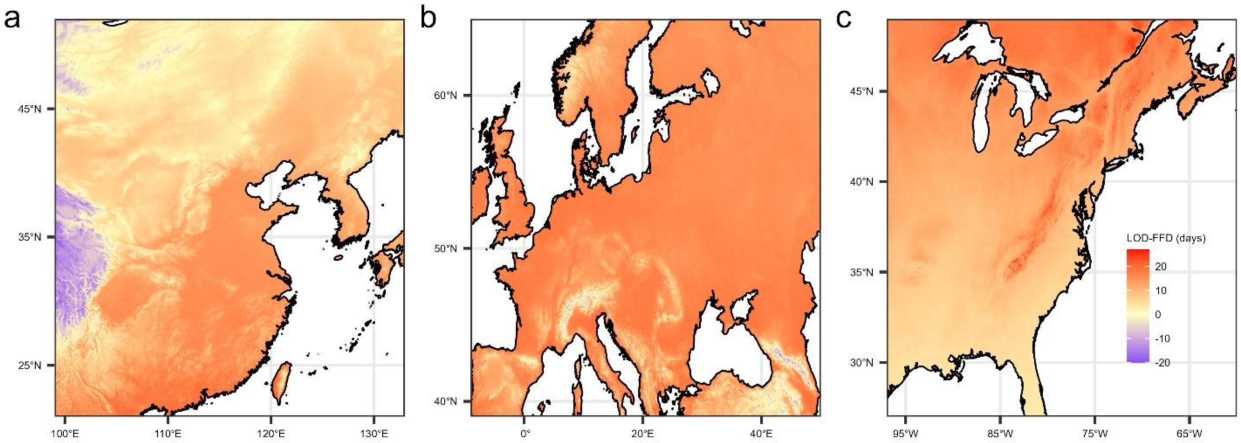

**Figure S12:** Full, uncropped version of Figure 4.

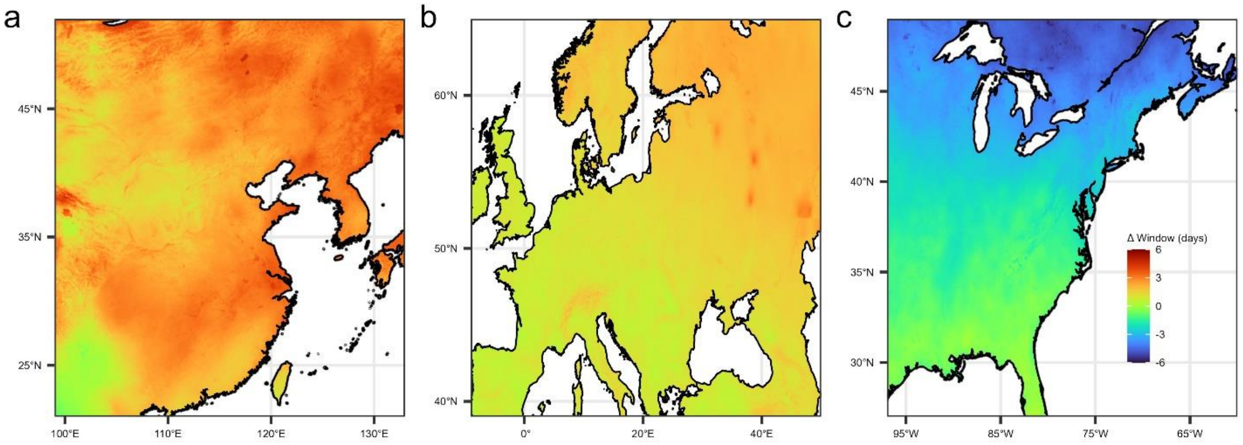

**Supplementary Information References:**

1. Ettinger, A. K. *et al.* Winter temperatures predominate in spring phenological responses to warming. *Nature Climate Change* **10**, 1137–1142 (2020).
2. Willems, F. M., Scheepens, J. F. & Bossdorf, O. Forest wildflowers bloom earlier as Europe warms: lessons from herbaria and spatial modelling. *New Phytologist* **235**, 52–65 (2022).
